# GEOAgent: An AI-driven Autonomous Framework for Intelligent GEO Data Retrieval and Standardized Preprocessing

**DOI:** 10.64898/2026.06.06.730646

**Authors:** Yingying Zhao, Quanyou Cai, Dongzhu Chen, Jiekai Chen

## Abstract

Datasets in the Gene Expression Omnibus (GEO) remain difficult to reuse at scale because sample annotations are heterogeneous and raw sequencing data require assay-specific preprocessing. We present GEOAgent, an AI-driven autonomous framework designed for intelligent dataset retrieval and standardized preprocessing by coupling autonomous semantic governance with an automated Nextflow pipeline named bioStream. Metadata from 181,760 sequencing series and 84,756 associated PubMed records were organized in a relational database and semantic index to support natural-language dataset retrieval. The framework automatically determines assay modalities, resolves experimental design pairings, and standardizes sample naming to minimize manual curation overhead. Based on these parsed attributes, the framework generates deployment-ready manifests to automatically execute containerized workflows across bulk and single-cell omics modalities. In expert-curated benchmarks, the workflow achieved 96% retrieval precision alongside 100% accuracy in assay classification and sample relationship resolution. The web platform is publicly accessible, while the source code and associated databases are openly available via GitHub and Zenodo.

## Introduction

The Gene Expression Omnibus (GEO) has served as a major public repository for functional genomics data since 2000^1,2^. As of April 2026, the database hosts over 281,000 public series comprising more than 8.4 million samples, providing a vast and invaluable resource for secondary research, meta-analyses, and data-driven scientific discovery. However, the full potential of this massive public asset remains heavily underutilized. Researchers attempting to reuse GEO datasets frequently encounter barriers caused by non-standardized sample descriptions, inconsistent annotations and assay-specific preprocessing requirements^3^. Previous studies have indicated that a substantial proportion of GEO submissions lack the structured annotations necessary for automated interpretation, necessitating extensive manual curation before any downstream computational analysis can begin^4^.

Furthermore, traditional keyword-based search interfaces often fail to bridge the semantic gap between imprecise natural language queries and the controlled vocabularies utilized in experimental documentation. Although compiled metadata repositories such as GEOmetadb have attempted to resolve querying inefficiencies by consolidating GEO attributes into a relational SQLite database, they fundamentally rely on rigid, exact SQL string matching^4^. Consequently, they remain highly vulnerable to nomenclature inconsistencies and cannot resolve implicit biological synonyms or acronyms. While existing platforms such as GEO2R, ARCHS4, and recount3 provide valuable web-based interfaces for differential expression analysis or offer pre-calculated expression matrices, they are constrained by inherent design limitations^5-7^. These platforms and static relational databases generally lack the architectural flexibility to process raw sequencing data through custom, standardized workflows, and they cannot autonomously interpret or decompose complex, multi-dimensional user intentions. Consequently, cross-study integration and large-scale data mining still demand substantial computational expertise (e.g., advanced SQL querying or pipeline engineering), leaving public genomics assets largely inaccessible to non-programming biologists.

To address these limitations and facilitate the reuse of public genomics data, we developed an integrated, open-source computational ecosystem comprising two core components: GEOAgent, an AI-driven intelligent retrieval and metadata curation platform, and bioStream, a containerized bioinformatic processing backend. Operating as a unified interactive assistant, GEOAgent leverages Large Language Models (LLMs) to bridge the gap between intuitive natural language intents and heavy data execution workflows^8^. The system implements a multi-stage Retrieval-Augmented Generation (RAG) architecture to resolve semantic ambiguities, seamlessly coupling a relational database with a dense vector store to guarantee precise semantic indexing and cross-encoder re-ranking^9^. Crucially, the framework moves beyond simple text mining by utilizing LLM-driven reasoning to automate complex metadata engineering tasks that typically require intensive manual intervention. GEOAgent autonomously conducts experimental modality and omics classification, standardizes sample identifiers and resolves sample relationships, including ChIP-seq input controls and single-cell Multiome (RNA/ATAC) pairing across disparate accessions. Once target cohorts are resolved, the fully curated metadata is dynamically handed off to bioStream, a standardized Nextflow backend configured with pre-built genome indices to coordinate highly reproducible, high-throughput multi-omics processing^10^. As of April 2026, the current implementation indexes 181,760 sequencing series from NCBI GEO and 84,756 linked PubMed publication records.

We evaluated this integrated workflow in three ways. First, we measure macro-dataset retrieval performance using 10 multi-constrained natural language queries and expert review of the top ten returned series for each query. Second, we assess the fine-grained accuracy of the platform’s cognitive auditing layer across three core metadata engineering tasks against manually reviewed ground-truth annotations^11^; this includes validating autonomous omics classification across 10 independent datasets, as well as resolving complex sample relationships across 10 paired single-cell multiome datasets and 10 independent ChIP-seq series to track antibody-to-input pairing fidelity. Third, we validated operational execution stability and pipeline scalability by testing the autonomous generation and end-to-end execution of bioStream preprocessing manifests on representative public datasets across multiple high-throughput assay categories. The resulting software implementation, indexed metadata assets, benchmark annotations, and containerized workflow configurations are fully released to support the open, reproducible reuse of public high-throughput sequencing data in alignment with FAIR principles^12^.

## Results

### Overview of the GEOAgent Platform for Automated High-Throughput Sequencing Data Reuse

To overcome the dual bottlenecks of metadata heterogeneity and preprocessing complexity in the Gene Expression Omnibus (GEO)^1,2,5^, we engineered GEOAgent, a modular, computational platform that supports high-throughput sequencing data curation, dissection, and workflow execution. The operational workflow and runtime architecture of GEOAgent are hierarchically organized into three functional layers spanning seven integrated processing steps, linking natural language user queries with reproducible bioinformatic workflows (Fig. 1).

**Figure 1.**
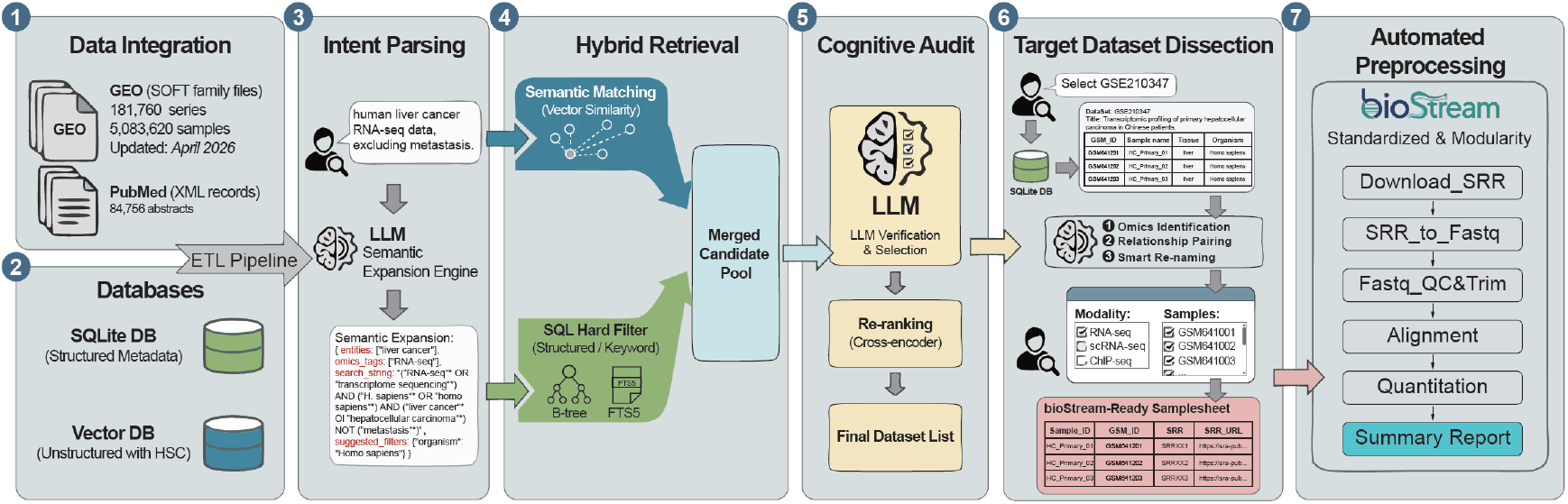
The schematic workflow of the GEOAgent. The end-to-end framework consists of seven interconnected modules. **(1) Data Integration:** Massive data from NCBI GEO (SOFT family files) and PubMed (XML records) are continuously ingested and processed via an automated ETL (Extract, Transform, Load) pipeline. **(2) Databases:** All integrated metadata are comprehensively stored and managed within a relational SQLite database. Concurrently, hierarchical semantic constructs (HSC) are converted into embeddings and stored in a Vector Database for similarity matching. **(3) Intent Parsing:** Natural language queries from users are parsed by a Large Language Model (LLM)-driven Semantic Expansion Engine to extract biological entities, omics tags, and construct optimized multi-clause search strings with suggested filters. **(4) Hybrid Retrieval:** The expanded query initiates a dual-path retrieval process, combining semantic matching (vector similarity) and SQL hard filtering (structured and keyword search), to yield a merged candidate dataset pool. **(5) Cognitive Audit:** An LLM-based verification and selection layer cross-examines the candidate datasets, followed by cross-encoder re-ranking to produce the final prioritized target list. **(6) Target Dataset Dissection:** The selected series (e.g., GSE210347) undergoes granular sample-level dissection, performing automatic omics type identification, relationship pairing, and smart sample re-naming to automatically generate a bioStream-ready samplesheet. **(7) Automated Preprocessing:** The structured samplesheet is streamed into *bioStream*, a modular Nextflow pipeline. This pipeline sequentially executes raw data downloading, format conversion, quality control, genomic alignment, and feature quantitation, generating a final summary report.

The infrastructure layer establishes a continuous data integration pipeline and a dual-engine storage backend. The framework ingests raw SOFT family files from GEO, encompassing 181,760 sequencing series and 5,083,620 individual samples, alongside 84,756 complementary XML literature abstracts from PubMed^13^ (Fig. 1, Step 1). Through an ETL (Extract, Transform, Load) pipeline, this heterogeneous multi-level metadata, which systematically integrates study, sample, platform, and corresponding literature layers, is decoupled and routed into a hybridized storage architecture designed to preserve both relational precision and biological context (Fig. 1, Step 2). Specifically, the comprehensive corpus of all retrieved metadata is structured and indexed within a centralized SQLite database, which is optimized with B-tree and FTS5 structures for robust relational querying and exact keyword filtering. To facilitate advanced semantic retrieval, the unstructured biological narratives within this master repository are concurrently processed via a specialized Hierarchical Semantic Chunking (HSC) strategy and converted into dense vector embeddings within a complementary vector database^14^.

This dual-engine backend directly fuels the system’s cognitive core, which functions as a multi-stage intelligent retrieval engine serving as the primary interactive interface for investigators. Upon receiving an open-ended natural language query, the framework establishes a bifurcated retrieval pipeline that processes the user intent through two distinct computational pathways (Fig. 1, Step 3). For the relational database branch, an LLM-driven Semantic Expansion Engine^15^ programmatically extracts biomedical entities and constructs boolean-bounded query constraints to capture complex relational variations. Conversely, for the vector database branch, the user’s original natural language query is dispatched directly to leverage the inherent semantic generalization capabilities of the dense vector space. These synchronized parallel branches subsequently converge in the Hybrid Retrieval pipeline, which combines the structured SQL keyword filtering with the semantic vector similarity matching to construct a merged candidate pool^16^ (Fig. 1, Step 4). To reduce false positive retrievals and guarantee high fidelity, this candidate pool is subjected to a rigorous Cognitive Audit layer (Fig. 1, Step 5). Within this auditing layer, deep cross-encoder re-ranking^17^ is coupled with an LLM-driven verification stage that evaluates candidates against strict biological exclusion criteria, producing a ranked list of candidate datasets.

The workflow culminates in the Target Dataset Dissection and Automated Preprocessing layers, which translate conceptual data requirements into reproducible, executable computational assets. Once a dataset of interest is selected, the agent instantly aggregates comprehensive study-level metadata and flattens the sample-level matrix into a unified view, reducing the need for manual navigation across GEO pages. Operating on this structured matrix, the agent automatically conducts fine-grained metadata engineering, including assay identification, sample renaming and complex experimental topology resolution such as aligning ChIP-seq input controls or pairing single-cell Multiome modalities (Fig. 1, Step 6). Instead of executing heavy computational workloads directly within the web interface, this automated curation process programmatically packages the resolved metadata into a deterministic, bioStream-ready samplesheet accompanied by pre-configured workflow execution scripts. This structured deployment package is designed to be downloaded or copied by investigators for local execution, ensuring full utilization of private high-performance computing infrastructure (Fig. 1, Step 7). The generated commands dynamically invoke bioStream, which functions as a standardized, containerized Nextflow backend^10^ deployed on the user’s local server or cluster environment. Guided by bioStream’s modular architecture, the standalone pipeline orchestrates a linear, fault-tolerant execution cascade spanning automated Sequence Read Archive (SRA) ingestion^18^, parallel quality control, genomic alignment, and feature quantification. This decoupled design ultimately delivers analysis-ready results and comprehensive summary reports to support downstream discovery while maintaining strict computational reproducibility and scalability.

### Knowledge Base Construction and Multi-Stage RAG Architecture for Precise Retrieval

The operational fidelity of the GEOAgent platform relies on a dual-database knowledge infrastructure designed to systematically resolve semantic ambiguities and execute complex queries. To construct this master repository, we compiled a comprehensive structured SQLite database that serves as the central data asset, housing 181,760 curated sequencing series, 5,083,620 individual samples, and 84,756 linked PubMed literature abstracts (Fig. 2). Relational analysis of this production instance reveals that the cohort is mainly composed of Homo sapiens and Mus musculus datasets, with RNA-seq serving as the predominant library strategy (Fig. 2B–D). However, further granular examination of these repository-wide statistics reveals metadata inconsistencies and indexing limitation in the public repository (Fig. 2C). While bulk RNA-seq represents the vast majority of labeled entries, standard library strategy metrics frequently fail to distinguish single-cell RNA-seq transcripts from bulk cohorts, while concurrently categorizing a massive proportion of datasets under a generic and uninformative ‘OTHER’ designation^4^. This annotation heterogeneity can reduce query precision in traditional search interfaces^12^. To mitigate this database-level limitation, our relational schema isolates the raw metadata into entity-specific tables comprising GSE metadata, GSM metadata, platform specifications, and PubMed records bound by explicit foreign-key mappings (Fig. 2A). Highly unstructured text fields are optimized using full-text search virtual tables to facilitate efficient keyword pinning, while categorical attributes are indexed via B-tree structures to accelerate multi-dimensional database slicing.

**Figure 2.**
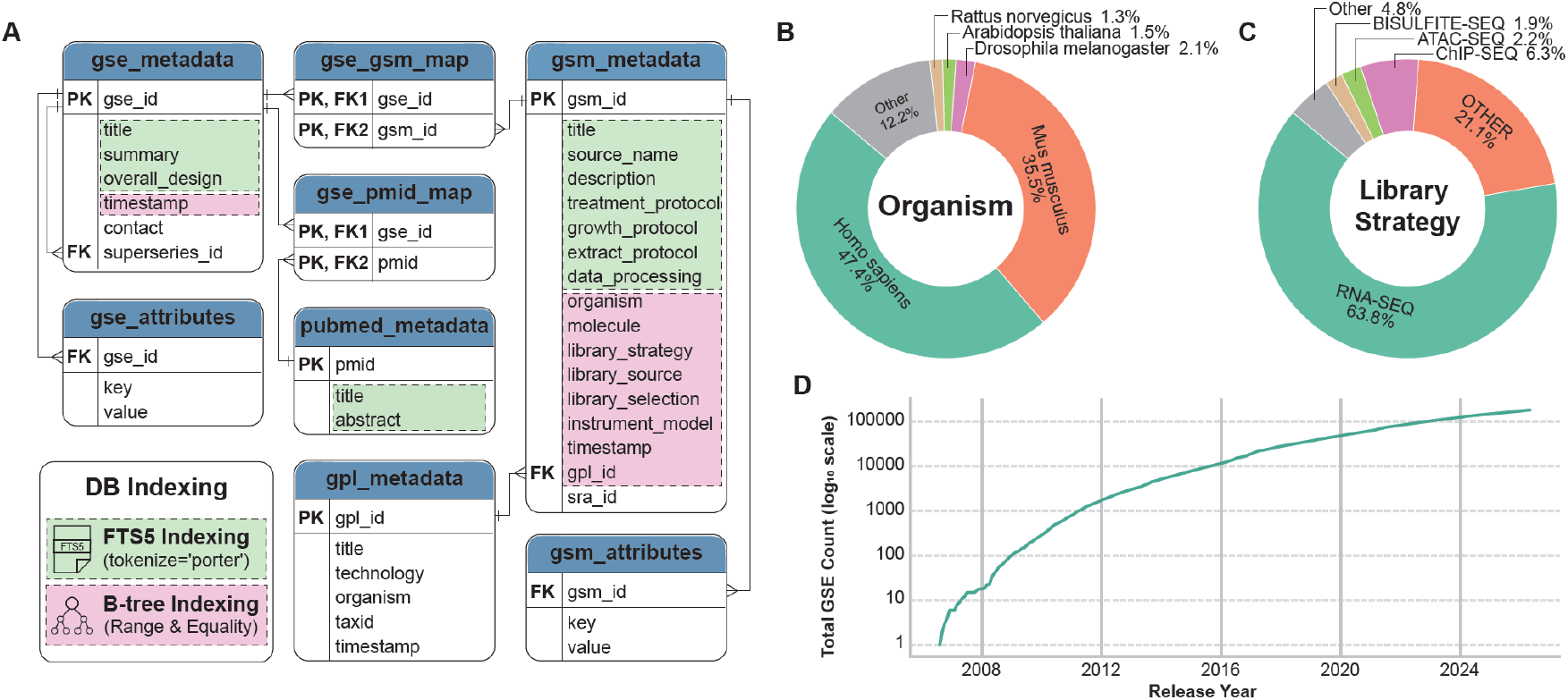
SQLite Database schema and data distribution of the integrated metadata. **(A)** Entity-relationship (ER) diagram of the local SQLite database. Green shading highlights text fields indexed via FTS5 (tokenize=‘porter’) for full-text search. Pink shading denotes fields optimized with B-tree indexing for range and equality queries. PK and FK represent primary keys and foreign keys, respectively. **(B)** Distribution of major biological organisms across the archived datasets, dominated by *Homo sapiens* and *Mus musculus*. **(C)** Composition of experimental data types based on library strategies, with RNA-SEQ representing the largest proportion. **(D)** Cumulative growth curve of the total GSE dataset count from 2008 to 2026.

To bridge the semantic gap deepened by these misannotation bottlenecks, the platform deploys a synchronized, dual-path parallel hybrid retrieval mechanism that concurrently drives relational and vector operations (Fig. 1, Step 4). This architecture processes incoming user queries through two asymmetric computational pathways to optimize candidate recall. For the relational database branch, the system utilizes an LLM-driven engine to decompose open-ended query intents, extract core biomedical nomenclature, and programmatically synthesize boolean-bounded search strings^19^. This expanded logical expression is routed exclusively to the SQLite full-text search index to keep explicit identifiers, gene symbols, and clinical terms in the candidate pool. Conversely, the vector database branch dispatches the user’s unexpanded original natural language expression directly into a dense vector database to capture fluid contextual similarities that bypass rigid text matching.

To populate this complementary vector repository without sacrificing the inherent experimental hierarchy of GEO records, we developed a Hierarchical Semantic Chunking (HSC) strategy (Fig. 3A). Rather than applying arbitrary character-length splitting, the metadata is partitioned based on explicit structural boundaries into discrete semantic zones, including core study profiles, sample descriptions, processing protocols, and associated publication abstracts (Table 1). These isolated text entities are then embedded into a high-dimensional space to form a semantic index.

**Table 1.**
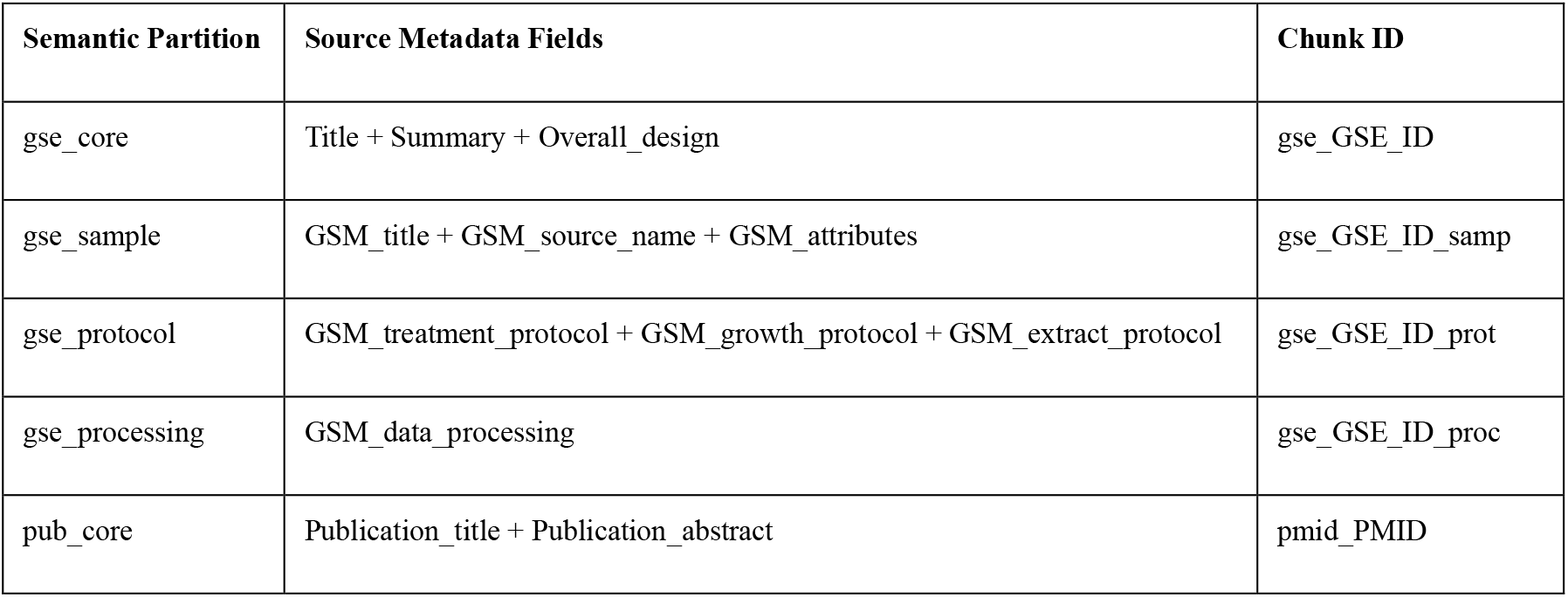
Semantic partitioning and chunk indexing schema for GEO and PubMed metadata. Columns define the structured partitions, their constituent source metadata fields extracted from NCBI records, and the standardized naming conventions used for downstream text embedding and vector retrieval (Chunk ID).

**Figure 3.**
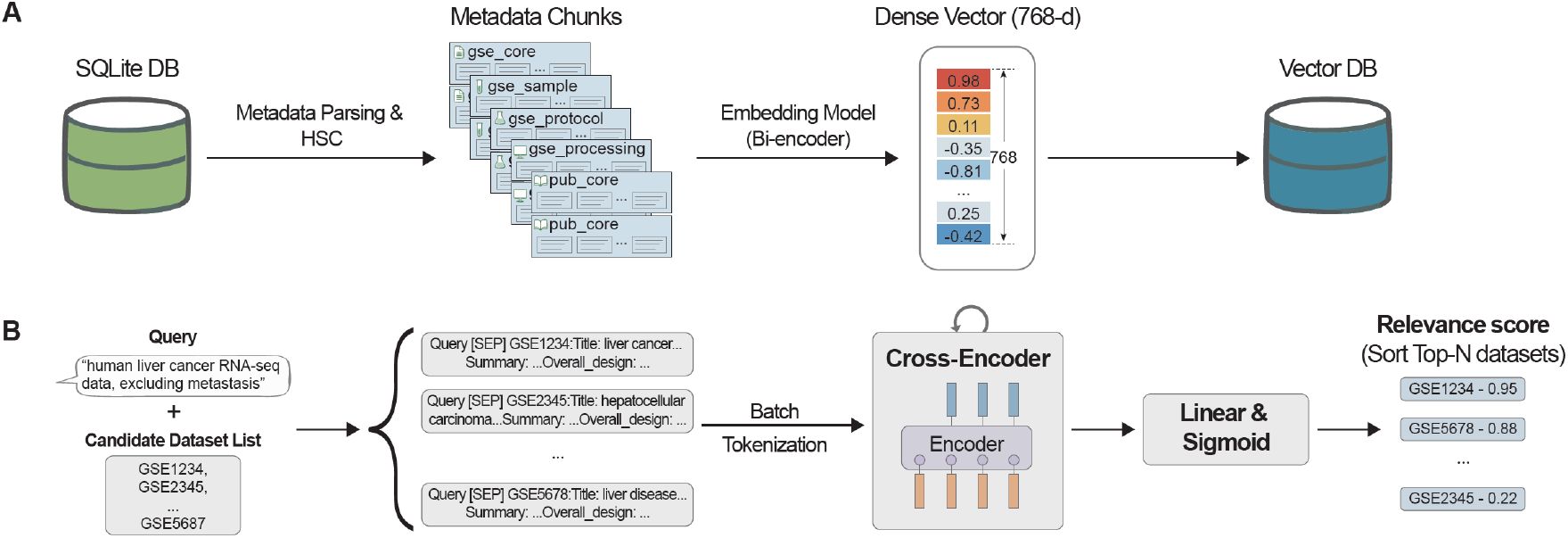
Workflow of vector database construction and dataset re-ranking. **(A)** Workflow for vector database indexing. Metadata from the SQLite DB are parsed into hierarchical semantic constructs (HSC) to generate metadata chunks (e.g., gse_core, gse_sample, gse_protocol). A bi-encoder embedding model converts these chunks into 768-dimensional dense vectors, which are subsequently stored in the Vector DB. **(B)** Pipeline for dataset re-ranking. The user query and the metadata of candidate datasets are concatenated using a [SEP] token and processed via batch tokenization. A Cross-Encoder performs deep attention-based encoding on the query-metadata pairs, followed by a linear layer and sigmoid activation to calculate individual relevance scores for sorting the top-N datasets.

To fine-tune the priority of the audited candidates, the remaining series are subsequently processed through a deep learning reranking pipeline (Fig. 3B). Within this scoring framework, the original user query and the retrieved metadata fields of each candidate dataset are concatenated into unified textual inputs. A deep Cross-Encoder^20^ then performs global attention-based encoding directly across these query-metadata pairs to map intricate contextual dependencies, effectively bypassing the limitations of shallow keyword matching. The resulting joint representations are utilized to compute cross-attended relevance scores, yielding a ranked list of GEO Series (GSE) candidates for the user query.

### Autonomous Metadata Engineering and Complex Topology Resolution

Beyond precise semantic retrieval, the core utility of the platform lies in its capacity to execute autonomous metadata engineering and complex topology resolutiosn, which traditionally bottleneck repository-scale data reuse due to their reliance on intensive manual curation (Fig. 1, Step 6). Public functional genomics datasets are frequently plagued by non-standardized nomenclatures, mislabeled assay modalities, and fragmented sample-to-sample relationships. To mitigate this data-governance challenge, the framework introduces an intelligent biological evaluation layer that programmatically unifies and standardizes unstructured text field descriptions, effectively transforming raw metadata noise into clean, analysis-ready operational structures.

This engineering pipeline initiates with a global assessment of assay modality to refine the coarse granularities and historical curation discrepancies observed across public records. Rather than relying on rigid metadata headers, which routinely lack the resolved vocabulary necessary to separate specialized genomic modalities from bulk cohorts or frequently collapse diverse sequencing techniques into an uninformative ‘OTHER’ designation, the platform cross-examines a comprehensive metadata ensemble encompassing study summaries, overall designs, and experimental protocols. By scanning for diagnostic technology footprints within these heterogeneous text blocks, the system accurately determines the precise omics type and library construction strategy of each dataset. Furthermore, this multidimensional parsing allows the framework to parse multi-omics datasets and assign standardized assay categories.

Once the underlying omics identities are verified, the framework dynamically resolves the alignment of complex experimental topologies, which are routinely fractured when researchers separate and upload paired biological samples across disjointed public accessions. By parsing unstructured experimental descriptions, the system maps the intricate relational links required for downstream comparative omics without human intervention. In chromatin immunoprecipitation sequencing (ChIP-seq) workflows, the system cross-references sample-specific phenotypic attributes with processing protocols to precisely pair target profiles with their corresponding baseline genomic input controls^21^. Similarly, in multi-omic studies where distinct cellular readouts from the same biological experiment are separated into disparate sample records, the framework reconstructs the underlying experimental design to align these disconnected sister datasets.

The resolution of these sample relationships subsequently enables a systematic normalization of heterogeneous sample nomenclature, removing a primary barrier to automated, parallel data consumption. Publicly deposited sample titles typically lack structured formatting or consistent delimiters, often varying substantially even within a single study. The platform reduces this naming inconsistency by extracting independent experimental variables, including treatment states, time-series intervals, genetic backgrounds, and biological replicate numbers, directly from the broader metadata environment. By synthesizing these isolated parameters, the framework successfully translates chaotic, highly idiosyncratic depositor names into clean, descriptive alphanumeric identifiers. This step produces structured metadata inputs for reproducible downstream workflow execution.

### End-to-End Automated Preprocessing and Multi-Omics Pipeline Execution

To ensure that the curated datasets can be immediately utilized for downstream biological discovery without imposing burdensome centralized infrastructure costs, the platform dynamically consolidates these verified omics designations, LLM-standardized sample nomenclatures, experimental topology mappings, and SRA download links accompanied by MD5 checksums into a unified configuration file tailored for bioStream (Fig. 1, Step 7). Operating as an industrialized localized framework managed through Nextflow^22^, this modular workflow orchestrates containerized multi-omics processing directly on the user’s independent computational infrastructure, systematically translating raw public sequencing entries into standardized, analysis-ready feature abundance matrices and genomic signal tracks (Fig. 4A).

**Figure 4.**
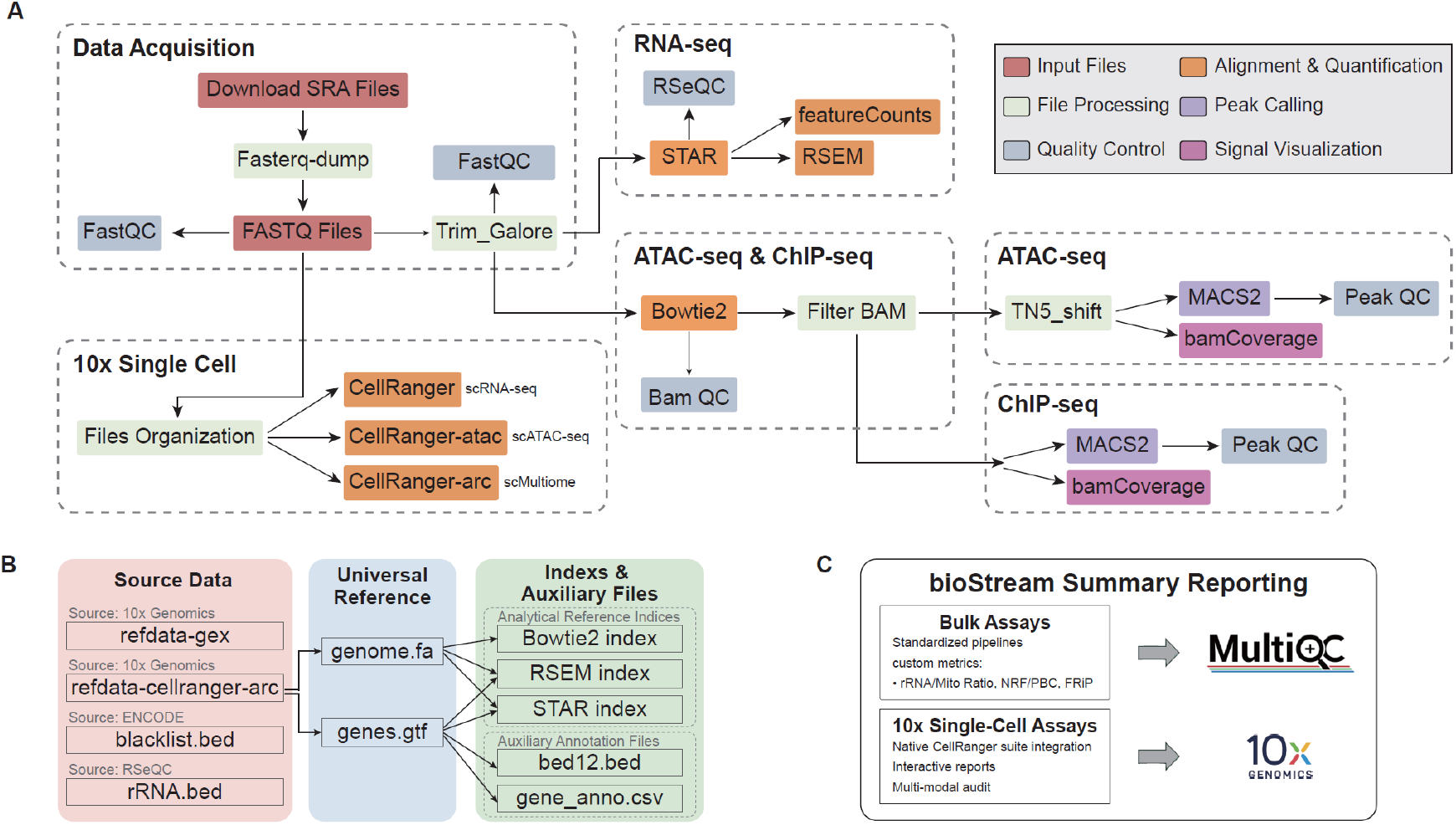
Overview of the bioStream processing pipeline. **(A)** Modular workflows for different assay modalities. Following data download and trimming, FASTQ files are routed to specific pipelines: RNA-seq (STAR, featureCounts, RSEM), ATAC-seq/ChIP-seq (Bowtie2, MACS2), or 10x Single Cell (CellRanger). Boxes are color-coded by operational tasks such as input, quality control, alignment, and peak calling. **(B)** Unified data sourcing and reference genome construction. Core reference files (genome.fa and genes.gtf) are sourced directly from 10x Genomics to build genomic indices for Bowtie2, RSEM, and STAR. These core files are also used to generate bed12.bed and gene_anno.csv as supporting files for the pipeline, alongside independent reference files from ENCODE (blacklist) and RSeQC (rRNA intervals). **(C)** Automated reporting and quality control aggregation. Bulk assays integrate native and custom-calculated metrics including rRNA/Mito ratios, NRF/PBC, and FRiP into a unified MultiQC report. For single-cell pipelines, native CellRanger summary reports are generated.

A primary technical challenge in cross-study integration is the reference genome heterogeneity that arises from disparate reference assemblies used across historical studies^23,24^. To reduce this source of variation during local deployment, the workflow implements a centralized genomic reference architecture anchored on a unified multi-ome reference framework (Fig. 4B). From this single unified reference source, the system programmatically derives the core genomic sequences and transcript annotations required to dynamically build identical downstream analytical indices, including STAR^25^, RSEM^26^, and Bowtie2^27^ models, while concurrently appending auxiliary genomic layout specifications such as ENCODE blacklist profiles^28^ and ribosomal RNA genomic intervals. This synchronized construction guarantees subsequent alignment and feature quantification steps within a consistent genomic coordinate space.

Leveraging these uniform genomic reference indices, the automated workflow routes sequence data through adaptive pathway branching based on the identified omics modalities (Fig. 4A). For bulk transcriptomics and epigenomics, raw sequence streams are processed through standardized alignment, peak-calling, or signal-quantification modules, followed by automated reporting and quality control aggregation (Fig. 4C). Crucially, the workflow extends standard reporting by integrating native and custom-calculated metrics, such as ribosomal RNA and mitochondrial contamination ratios (rRNA/Mito ratios), library complexity indices (NRF/PBC), and fraction of reads in peaks (FRiP)^21^, into a unified MultiQC report^29^. Conversely, for 10x Genomics single-cell assays, the workflow directs data through corresponding CellRanger suites to generate native summary reports. By combining customized metric aggregation for bulk datasets with native 10x Genomics summary outputs, this highly portable workflow enables users to execute fully automated raw SRA data downloading and multi-omics preprocessing on local computing systems.

### Implementation of the GEOAgent Graphical User Interface and End-to-End Workflow Validation

To demonstrate the practical utility, accessibility, and engineering robustness of the platform, we developed a web-based graphical user interface (GUI) for GEOAgent, providing investigators with an intuitive and transparent data-mining workspace (Fig. 5). The runtime architecture, automated data governance, and modular execution pipeline of the underlying framework were validated through an end-to-end operational case study focusing on public liver cancer sequencing cohorts.

**Figure 5.**
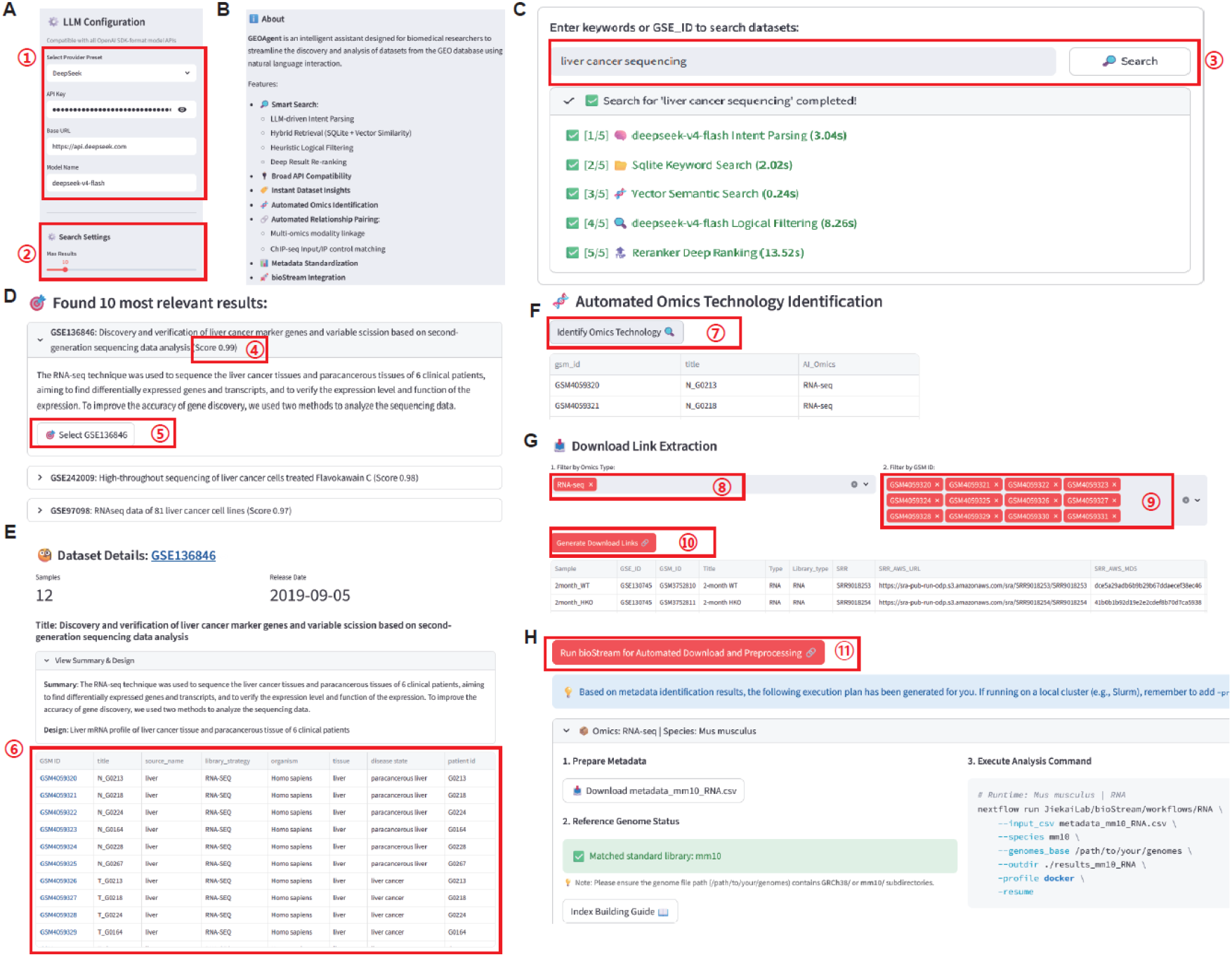
User interface components and interactive search workflow of GEOAgent. **(A)** LLM configuration and search setting panels. Red boxes 1 and 2 highlight the API provider selection (e.g., DeepSeek) and the configuration of search result limits, respectively. **(B)** Overview of GEOAgent functional features and system characteristics. **(C)** Natural language search input and multi-stage backend execution trace. Box 3 shows the query interface, with logs capturing the response times for intent parsing, keyword search, vector search, LLM filtering, and ranking. **(D)** Prioritized dataset search results. Box 4 indicates the computed relevance score, and box 5 provides the action button to select a target series (e.g., GSE136846). **(E)** Sample-level metadata extraction page for the selected series. Box 6 highlights the dynamic visualization panel, which automatically retrieves and displays the full set of structured sample attributes (e.g., title, source name, organism, tissue, and dataset-specific covariates) provided by the original GEO record. **(F)** Automated omics technology identification module. Box 7 triggers the automatic classification of sample-level assay modalities. **(G)** Target filtering and selection for sequencing data download. Boxes 8 and 9 manage the filtering and selection of targeted omics types and individual samples of interest, respectively. Box 10 executes the backend parsing logic, which automatically performs sample re-naming, secondary omics pairing for single-cell multi-omics, and Input-IP control matching for ChIP-seq assays, prior to generating the download links. **(H)** Execution interface for downstream pipelines. Box 11 initiates the automated data downloading and preprocessing via the BioStream framework, providing structured metadata downloads and reproducible execution commands.

The control dashboard provides flexible environment initialization, supporting the direct configuration of large language model deployment via an OpenAI SDK-compatible API (Fig. 5A, ①) and customizable search depth thresholds to balance retrieval latency against metadata catalog coverage (Fig. 5A, ②). Upon receiving an open-ended, natural language query such as “liver cancer sequencing” (Fig. 5C, ③), the retrieval engine executes a multi-stage search strategy. The interface explicitly renders a real-time agentic reasoning log capturing the exact execution latencies for intent parsing, structured SQLite keyword queries, vector semantic matching, logical filtering, and cross-encoder re-ranking, ultimately returning the top 10 candidate series ranked by their structural similarity scores (Fig. 5C-D, ④).

Selecting a high-scoring candidate cohort (e.g., GSE136846) dynamically populates the dataset dissection workspace (Fig. 5D-E, ⑤). Rather than requiring manual navigation across disjointed repository pages, the interface aggregates cross-layered study attributes into a unified, high-density dashboard that exposes the project title, experimental summary, and complete sample-level metadata matrices (Fig. 5E, ⑥). Activating the automated technology identification controller (Fig. 5F, ⑦) triggers the agentic layer to audit sample entries and determine true empirical assay types, standardizing heterogeneous depositor names into definitive computational modalities (e.g., RNA-seq).

Following cohort selection and filtering, users can extract download links tailored to specific omics classifications and sample subsets (Fig. 5G, ⑧-⑩). During this transition, the platform programmatically links individual GSM entries to their corresponding Sequence Read Archive (SRA) source paths, generating a standardized data manifest containing cloud storage URLs and MD5 checksums.Finally, this structured manifest is integrated into the workflow invocation panel, where the workflow assigns the standard reference assembly based on the cohort’s species (Fig. 5H). The dedicated bioStream execution controller (Fig. 5H, ⑪) synthesizes a compiled Nextflow command string containerized via Docker. This closed-loop design eliminates manual environment engineering and programming overhead, allowing researchers to transition from unstructured natural language queries to reproducible pipeline execution within a single interface.

### Quantitative Benchmarking of Retrieval Accuracy and Curation Efficiency

While the graphical interface simplifies user interaction, quantifying the objective retrieval fidelity and computational efficiency of the platform is essential. Therefore, we evaluated the platform’s performance using a probe-based validation strategy across ten distinct genomic search scenarios designed as benchmark queries (Fig. 6). Given the heterogeneous and expanding nature of public databases, establishing an exhaustive global ground truth is practically unfeasible. To circumvent this baseline limitation, expert manual curation was employed to assess the biological relevance of the retrieved results under a maximum return threshold of 10 entries per scenario (Fig. 6A; detailed in Supplementary Table S1). Across all ten complex query scenarios, the platform achieved an overall retrieval precision of 96%, successfully identifying 47 true positive dataset series out of 49 total retrieved entries (Fig. 6A).

**Figure 6.**
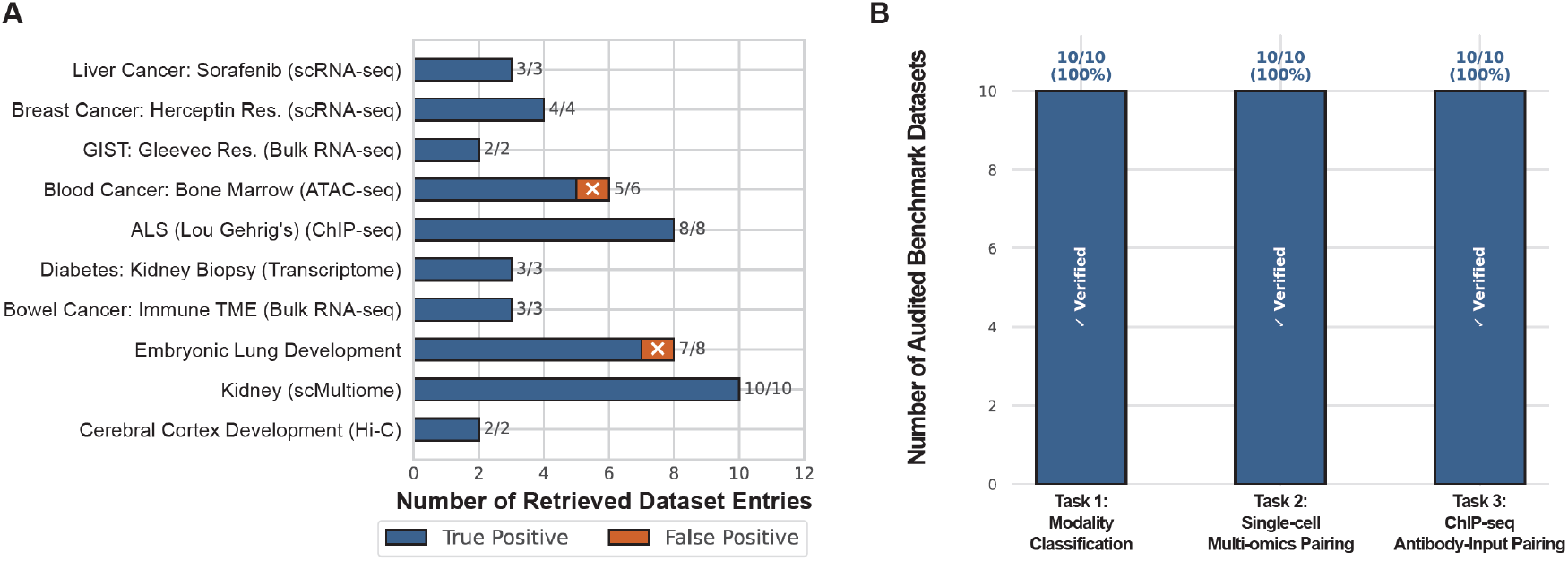
Expert-curated benchmark validation of GEOAgent performance. **(A)** Retrieval accuracy across ten complex biomedical query scenarios under a maximum result threshold of 10. True positive (blue) and false positive (orange) classifications for all retrieved entries were mutually verified through expert manual curation. **(B)** Expert review of three core analytical tasks evaluated on 10 benchmark datasets. Manual verification confirmed a 100% (10/10) accuracy rate for GEOAgent in assay modality classification (Task 1), single-cell multi-omics relationship pairing (Task 2), and ChIP-seq antibody-input control matching (Task 3).

Compared to traditional functional genomics cataloging utilities such as GEOmetadb, which rely on rigid relational database architectures (SQLite) and require advanced SQL querying and exact syntax, the platform implements a hybrid dual engine that couples relational querying with vector semantic databases to tolerate natural language queries and interpret open-ended search intents (Table 2). This architecture reduces the burden of manual query construction and adds tolerance for semantic variation. Consequently, the platform’s retrieval benchmark demonstrated high precision across diverse biomedical and pharmacological semantics, successfully resolving nomenclature heterogeneities and clinical-to-technical phrase variations without literal string matching (Fig. 6A, Supplementary Table S1). For example, in the query “liver cancer scRNA-seq sorafenib treatment”, the system bypassed raw text restrictions to retrieve series where the disease was specified as “hepatocellular carcinoma” and the modality was cataloged as “Single-cell RNA sequencing”.

**Table 2.**
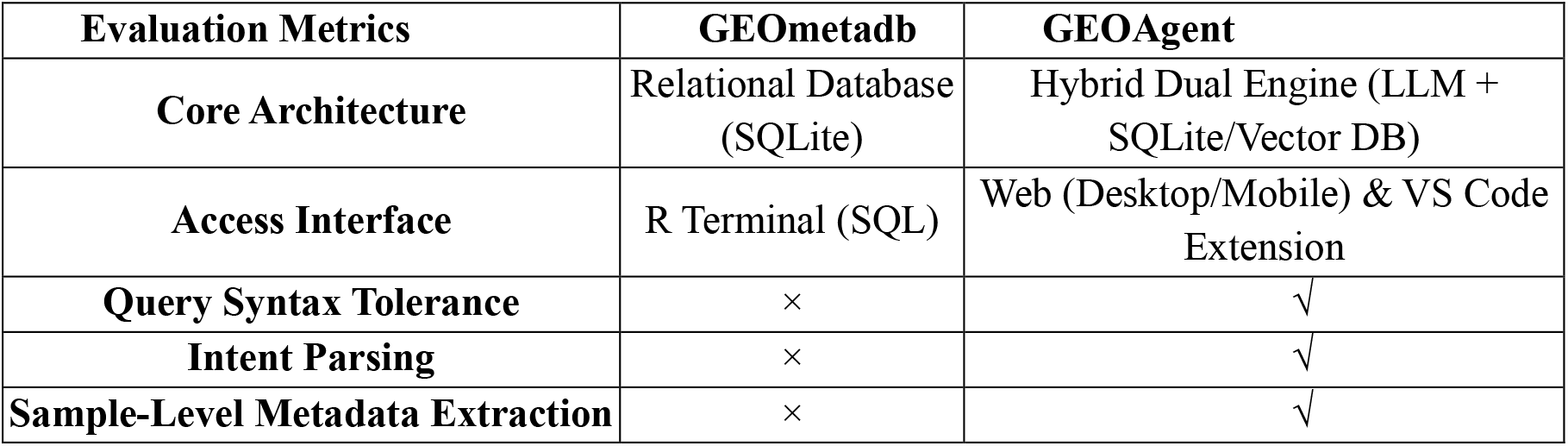
Feature and architectural comparison between GEOmetadb and GEOAgent. Rows itemize critical technical benchmarks including core database architecture, user access entry points, natural language tolerance, search intent interpretation, and deep sample-level metadata processing capabilities.

| Evaluation Metrics | GEOmetadb | GEOAgent |
| --- | --- | --- |
| Core Architecture | Relational Database (SQLite) | Hybrid Dual Engine (LLM + SQLite/Vector DB) |
| Access Interface | R Terminal (SQL) | Web (Desktop/Mobile) & VS Code Extension |
| Query Syntax Tolerance | × | √ |
| Intent Parsing | × | √ |
| Sample-Level Metadata Extraction | × | √ |

Furthermore, the framework demonstrated robust generalization from commercial drug trade names to generic counterparts and precise biological mechanisms. In stress tests using the query “scRNA-seq datasets evaluating Herceptin resistance in breast cancer”, the framework automatically expanded the commercial token to its target mechanism, accurately retrieving series annotated with “HER2-targeted therapies” or “dual ER/HER2 blockade”^30^. Similarly, for the query targeting “Gleevec resistance in gastrointestinal stromal tumors (GIST)”, where the literal term “Gleevec” was entirely absent from the target repository metadata, the platform successfully matched the underlying synonymy with the generic compound Imatinib, deeply parsing dataset summaries to identify active profiling of sensitive versus resistant cohorts(Fig. 6A, Supplementary Table S1).

Beyond text retrieval, the efficiency of the platform’s downstream metadata processing was validated through expert review of three core curation tasks across 10 dedicated benchmark datasets per task, totaling 30 audited dataset instances (Fig. 6B). In contrast to conventional methods that lack sample-level metadata extraction capabilities (Table 2), the platform completed these cognitive auditing tasks within seconds, achieving a 100% (10/10) accuracy rate across all evaluated categories (Fig. 6B). For Task 1 (Assay modality classification), the system correctly identified underlying sequencing technologies and library strategies from non-standardized text. For Task 2 (Single-cell multi-omics relationship pairing) and Task 3 (ChIP-seq antibody-input control matching), the parsing logic successfully reconstructed the original experimental designs by linking companion single-cell modalities and pairing biological immunoprecipitation tracks with their corresponding genomic input controls. These benchmarks demonstrate that the framework significantly minimizes the manual curation burden while maintaining expert-level accuracy, providing a scalable solution for high-throughput genomic data reuse.

## Discussion

The integration and reuse of public functional genomics datasets are important forbiomedical research, but they remain limited by heterogeneous depositor-submitted metadata^31^. While repositories such as NCBI GEO and SRA host large volumes of multi-omics data, the lack of standardized nomenclature, unstructured sample annotations, and mislabeled assay modalities frequently make automated reuse difficult. Traditional methods to address this metadata problem typically rely on manual expert curation, which is inherently unscalable, or rigid keyword matching systems that fail to resolve biological synonyms or experimental topographies. In this study, GEOAgent shows that LLM-assisted parsing, when combined with rule-based checks and expert-curated prompts, can support metadata standardization for the evaluated tasks. By decoupling unstructured textual descriptions from the downstream analytical configuration, the platform effectively converts heterogeneous public metadata into structured records for downstream workflow execution.

A key objective of this study was to reduce the practical barrier to high-throughput bioinformatics workflows for experimental biologists and clinical investigators. Historically, leveraging structured repository metadata often required computational expertise, such as executing complex multi-table joins and exact regular expressions within relational database environments like GEOmetadb^4^. This technical barrier has limited independent use of public data cohorts by many experimental researchers. The interface reduces this burden by wrapping database queries, Nextflow workflow configuration and Docker-based execution in a natural-language workflow. By generating execution commands that incorporate SRA paths and reference assemblies, the framework allows users to initiate local multi-omics preprocessing with less manual command-line setup.

Despite these advancements, several technical limitations must be acknowledged to guide future deployment. First, although the underlying engine implements localized rule-based constraints and task-specific prompts to govern model outputs, large language models can still produce variable or incorrect outputs^32^. This risk is likely higher for uncommon compound names, recently introduced assays or metadata patterns that are poorly represented in model training data. Second, while the preprocessing workflows support standard bulk omics (RNA-seq, ATAC-seq, and ChIP-seq) and 10x Genomics single-cell assays, the pipeline architecture does not yet natively accommodate alternative high-throughput modalities. Future updates will expand the platform’s processing spectrum to support a wider array of bulk assays including Hi-C and DNA methylation sequencing, as well as non-10x single-cell platforms such as plate-based Smart-seq2 and emerging spatial transcriptomics technologies without requiring manual downstream intervention. Finally, while the web-based graphical interface has low centralized computational costs, the execution efficiency of the generated bioStream pipelines remains bounded by the user’s local hardware constraints and cluster architecture configurations.

Future development will focus on broader assay compatibility and optional downstream analysis modules. Currently, bioStream is engineered to streamline upstream data preprocessing, specifically focusing on standardized raw sequence ingestion, sequence alignment, feature quantification, and quality control. Future versions will establish an autonomous downstream analytical framework to automate higher-level cognitive tasks, including cell-type annotation, differential expression analysis, and functional pathway enrichment. Because downstream analysis depends on study context and hypothesis testing, these modules will require additional validation. To support these operations, we aim to integrate specialized biomedical foundation models and domain-specific knowledge graphs, such as MeSH and UMLS^33,34^, directly into the platform to provide controlled vocabulary support and reduce ambiguous entity matching. Concurrently, the upstream pipeline will be updated to incorporate containerized workflows for alternative single-cell modalities, long-read sequencing, and spatial genomics platforms. Ultimately, we envision this system evolving into a fully autonomous data-driven ecosystem, enabling researchers to seamlessly transition from a baseline biological hypothesis expressed in natural language to a thoroughly curated, processed, and downstream-interpreted multi-omics data matrix and its associated biological insights.

## Methods

### 1. Data Ingestion and Structured SQLite Database Construction

#### 1.1 Data Acquisition and Inclusion Criteria

Metadata synchronization was performed using the NCBI Gene Expression Omnibus (GEO) FTP server^5^. The inclusion criteria strictly targeted datasets utilizing high-throughput sequencing technologies, while legacy microarray-only studies were excluded. Dataset accessibility was verified by the presence of SOFT-formatted family files. Datasets with corrupted formatting or extraction failures were programmatically excluded. To enrich the repository with bibliographic evidence, the validated series were linked to the NCBI PubMed database via the Entrez Utilities API to retrieve the titles and abstracts of study-associated publications^35^. These assets were systematically synchronized into both the relational and vector databases, establishing a dual-source knowledge base linking literature-level narratives with primary omics records.

### 1.2 Relational Schema Design and Table Architecture

The SQLite database structure was determined empirically by scanning the statistical coverage of all metadata keys across the retrieved GEO SOFT files (Fig. 2A). A field was designated as a standalone relational column only if it demonstrated near-100% historical prevalence and held essential biological relevance. High-coverage fields were partitioned into specialized core tables: gse_metadata for series-level summaries, gsm_metadata for sample records, and gpl_metadata for platform technologies.

To prevent table fragmentation caused by highly variable and study-specific descriptions, heterogeneous experimental variables and dynamic annotations were decoupled from the core schema. Attributes that fluctuate drastically between distinct datasets—ranging from clinical phenotypes (e.g., patient age or tissue pathology) to customized assay conditions (e.g., treatment doses, time points, or genetic perturbations)—were programmatically routed into long-format Entity-Attribute-Value (EAV)^36^ tables (gse_attributes and gsm_attributes). This layout ensures infinite horizontal scalability for highly sparse annotations while maintaining a clean database core.

For literature linking, unique PubMed IDs extracted from the GSE entries were mapped directly to a dedicated pubmed_metadata table through a standard junction table (gse_pmid_map). Complex many-to-many topologies between series and individual samples were natively resolved using an identical relational mapping strategy via gse_gsm_map.

### 1.3 Database Indexing Strategy and Full-Text Search (FTS5)

To ensure sub-second query performance across the relational data layer, a dual-indexing strategy was deployed within the SQLite engine to balance unstructured text parsing against structured categorical filtering (Fig. 2A):

1. Full-Text Search (FTS5) Indexing: For highly unstructured text fields, FTS5 virtual tables were constructed using the Porter stemming algorithm^37^. This tokenizer programmatically reduces words to their baseline morphological roots (e.g., stemming “sequenced”, “sequencing”, and “sequences” to “sequenc”). This virtual index encapsulates multi-level free-text attributes, including study scopes (gse_metadata: title, summary, and overall_design), sample processing descriptions (gsm_metadata: title, source_name, description, treatment_protocol, growth_protocol, extract_protocol, and data_processing), and literature abstracts (pubmed_metadata: title and abstract). This index native supports rapid prefix wildcard (*) and Boolean logical queries during downstream database pruning.
2. B-tree Indexing: For controlled vocabularies and temporal constraints, standard B-tree indexes were mapped to rigid categorical columns^38^. These include fixed metadata fields within the gsm_metadata table—such as organism, molecule, library_strategy, library_source, library_selection, and instrument_model—as well as chronological registration timestamps across all series and sample tables. This hybrid indexing topology guarantees O(log n) time complexity for exact match filtering, categorical aggregation, and range-based temporal queries, providing the structural performance required for real-time iterative data filtering.

### 2. Metadata Semantic Chunking and Vector Database Construction

#### 2.1 Hierarchical Semantic Chunking (HSC)

To transform unstructured metadata into semantically cohesive knowledge blocks without fragmenting experimental context, a structure-aware Hierarchical Semantic Chunking (HSC) pipeline was implemented (Fig. 3A). Instead of applying fixed-token-length splitting^39^, the HSC framework partitions metadata based on the logical boundaries of GEO and PubMed entities. Metadata fields are programmatically aggregated into five distinct semantic partitions, as detailed in Table 1: gse_core, gse_sample, gse_protocol, gse_processing, and pub_core.

To prevent vector database bloating and eliminate text redundancy caused by identical protocol descriptions shared across multiple samples within a single dataset, an intra-series de-duplication workflow was deployed for the gse_protocol and gse_processing partitions. Identical text strings describing growth, treatment, or computational processing workflows across different samples of the same study are collapsed into a singular, unique knowledge block linked back to the parent series ID.

To manage exceptionally dense descriptive text, the HSC architecture enforces an upper size ceiling at a maximum threshold of 2,000 tokens per block. If a consolidated metadata field breaches this 2,000-token limit, a symbolic backoff mechanism triggers a recursive split. This fallback workflow sub-tokenizes the text along natural linguistic delimiters, prioritizing paragraph breaks (\n), sentence periods (.), and semicolons (;) to maintain localized contextual coherence.

#### 2.2 Vector Space Mapping and Storage

Following chunk optimization, each discrete metadata block is mapped into a continuous vector space to enable semantic similarity retrieval. Textual chunks are vectorized using the Nomic Embed v2 embedding model^40^, which generates 768-dimensional dense vectors (Fig. 3A). To ensure data privacy and eliminate runtime API latencies, the embedding network is hosted locally within the computing infrastructure via the Ollama orchestration engine. The resulting multi-dimensional dense embeddings are subsequently ingested and indexed within a localized vector storage layer, supporting efficient semantic similarity queries.

### 3. Multi-Stage Hybrid Retrieval and Cross-Encoder Re-Ranking Pipeline

#### 3.1 Multi-Stage Hybrid Retrieval Architecture

The runtime retrieval engine executes a sequential multi-stage workflow to optimize candidate dataset recall and precision (Fig. 1). The pipeline progresses through intent parsing, parallel hybrid retrieval, LLM-driven screening, and deep re-ranking:

1. **Intent Parsing and Semantic Expansion:** Upon receiving a natural language query, a local large language model (LLM) extracts core biomedical entities—such as genes, diseases, and cell types—and expands them with scientific synonyms. This entity payload is programmatically compiled into a structured JSON dictionary and a Boolean logic query string optimized with wildcard (*) modifiers.
2. **Dual-Path Parallel Hybrid Retrieval:** The search is executed simultaneously across both data layers. The expanded Boolean string runs as a hard filter across the SQLite FTS5 index for precise keyword pinning. Concurrently, the raw natural language query is routed to the vector database to compute cosine similarity scores across the dense embeddings. Candidate series from both pathways are merged during this phase.
3. **LLM-Driven Exclusion Screening:** The merged candidate cohort is subjected to an LLM-driven filtering layer operating on strict rule-based exclusion prompts. The model evaluates the unstructured metadata chunks to purge false positives. Candidates with mismatched sequencing modalities (e.g., bulk RNA-seq when single-cell RNA-seq was requested), discordant biological systems (e.g., in vitro organoids when native tissues were requested), or unaligned treatment conditions are programmatically excluded.

#### 3.3 Deep Cross-Encoder Re-Ranking

Surviving datasets from the initial screening are passed through a deep re-ranking pipeline utilizing the bge-reranker-v2-m3 Cross-Encoder model to determine final priority (Fig. 3B). Unlike bi-encoders that evaluate queries and documents independently^20^, the Cross-Encoder architecture forces the user query and the dataset text to interact directly via full joint-attention mechanisms.

In practice, the system pairs the user query with each candidate dataset. For each candidate, its core metadata fields—comprising the title, summary, and overall design—are concatenated into a singular text block. The query and the metadata block are then joined using a standard separation token ([SEP]). To optimize computational throughput, these text pairs are processed via a batch tokenization layer to generate standardized input tensors.

These tensors are funneled into the Cross-Encoder network, which performs token-to-token attention mapping between the query constraints and the dataset descriptions. The resulting encoder output is passed through a linear and sigmoid activation layer to calculate a normalized relevance score bounded between 0 and 1. The candidate datasets are then sorted in descending order based on these relevance scores to yield the finalized priority list of target genomics series for downstream execution.

### 4. Automated Metadata Curation and Experimental Topology Resolution

Prior to downstream workflow orchestration, the framework executes automated metadata transformations and topology resolution via an LLM-driven curation layer (Fig. 1, Block 6). This pipeline standardizes unstructured annotations and maps experimental relationships through three functional modules:

1. **Omics Identification:** To correct inconsistencies and mislabeling in public metadata submissions, an automated evaluation protocol is implemented to classify the exact omics modalities and assay technologies of individual sample accessions. The curation engine is fed a comprehensive metadata package encompassing global series contexts, tabular phenotype attributes, and de-duplicated protocol strings. By enforcing a single-cell priority heuristic and scanning for technology-specific syntax (e.g., 10x Chromium, Smart-seq2, or Cell Ranger outputs), the system differentiates between conventional bulk assays and single-cell platforms. Crucially, the workflow resolves multi-modal co-assays (e.g., separating dual-modality single-cell Multiome from isolated single-modality scRNA-seq or scATAC-seq runs) and programmatically outputs a unified classification tag following standardized nomenclature.
2. **Relationship Pairing:** To eliminate manual annotation overhead and streamline downstream data reuse, the framework parses unstructured text strings to map complex sample topologies. The engine programmatically identifies and pairs corresponding control and experimental designs without manual intervention. This includes automatically coupling chromatin immunoprecipitation sequencing (ChIP-seq) input files with their respective immunoprecipitation (IP) samples^21^, as well as aligning split single-cell multi-omics partner datasets (e.g., paired RNA-seq and ATAC-seq fractions) that were uploaded across separate GSM accessions.
3. **Smart Re-naming:** To eliminate laborious manual mapping and facilitate direct programmatic parsing, a rule-based generative schema is deployed to standardize sample identifiers. Because raw GSM accessions lack biological context and original sample titles are frequently non-standardized or excessively redundant, the system evaluates localized metadata variables—such as treatment conditions, time points, and biological replicates. These key variables are programmatically synthesized to automatically rename individual records into clean, descriptive, and delimited identifier strings, eliminating the need for manual curation during batch downstream analysis.

### 5. Workflow Orchestration and Standardized Preprocessing Pipelines

#### 5.1 Nextflow Framework and Reference Standardization

After cohort selection and metadata curation, GEOAgent exports a standardized samplesheet containing sample identifiers, curated assay labels, species information, SRA links, and inferred pairing relationships. This manifest is consumed by bioStream, a Nextflow-based workflow^10^ that executes containerized modules through Docker or Singularity on local workstations or HPC environments. To preserve cross-study comparability, bioStream uses a standardized reference configuration derived from the 10x Genomics refdata-cellranger-arc resource and supplemented with ENCODE blacklist^28^ and RSeQC rRNA annotations^41^. Detailed command-level execution, local deployment, and reference-building procedures are provided in Supplementary Note 1.

#### 5.2 Multi-Omics Processing Modules

1) Assay-specific workflow branches were selected from the GEOAgent-curated modality tags. Bulk RNA-seq datasets were processed through STAR-based alignment^25^ and transcript or gene-level quantification; ATAC-seq and ChIP-seq datasets were processed through Bowtie2-based alignment^27^, BAM filtering, peak calling, and signal-track generation; and 10x Genomics scRNA-seq, scATAC-seq, and single-cell Multiome datasets were processed through the corresponding Cell Ranger workflows. The main manuscript summarizes the analytical design and outputs, whereas tool-level execution details are retained in Supplementary Note 1.

#### 5.3 Automated Quality Auditing and Unified Reporting

Quality-control outputs were aggregated into assay-appropriate reporting layers. Bulk RNA-seq, ATAC-seq, and ChIP-seq workflows were summarized through MultiQC^29^ together with custom metrics such as rRNA/Mito ratios, NRF/PBC, and FRiP^21^, whereas 10x Genomics workflows retained native Cell Ranger web summaries. This reporting strategy provides a unified audit layer while keeping assay-specific quality indicators interpretable for downstream reuse; implementation-level reporting details are provided in Supplementary Note 1.

### 6. Evaluation Framework and Expert-Curated Benchmarks

#### 6.1 Retrieval Performance Evaluation

A benchmark framework was constructed to evaluate the retrieval and metadata-curation capabilities of GEOAgent. The retrieval benchmark comprised 10 distinct natural-language queries designed to span a broad spectrum of biological and technical use cases—including disease-specific bulk RNA-seq searches, single-cell queries, epigenomic assays, strict exclusion criteria, and multi-modal study designs (Fig. 6A). All queries were locked prior to model evaluation; query string configurations and their detailed evaluation criteria are documented in Supplementary Table S1. To rigorously test the platform’s re-ranking and logical filtering capacity under high-density constraints, the maximum return limit for each query was strictly capped at the top 10 dataset entries.

For retrieval auditing, two domain experts independently reviewed all candidate Gene Expression Omnibus (GEO) series returned within the top 10 results to assign binary relevance labels (True Positive or False Positive) according to predefined inclusion and exclusion criteria. Discrepancies between the two primary reviewers were systematically resolved and finalized under the adjudication of a third senior reviewer to guarantee absolute annotation reliability.

#### 6.2 Metadata Curation Evaluation

A separate curation benchmark was established to evaluate the three core metadata-parsing tasks, utilizing a total of 30 expert-annotated GEO series (10 unique datasets dedicated per task; Fig. 6B):

1. **Task 1 (Omics Identification):** Classifying exact assay platforms or sequencing chemistry configurations from highly heterogeneous text metadata.
2. **Task 2 (Single-cell multi-omics relationship pairing):** Programmatically linking corresponding single-cell multi-omics fractions (specifically matching gene expression [GEX] and chromatin accessibility [ATAC] files) originating from identical biological samples but split across separate GSM accessions.
3. **Task 3 (ChIP-seq antibody-input control matching):** Correctly pairing biological chromatin immunoprecipitation sequencing (ChIP-seq) immunoprecipitation (IP) samples with their respective antibody-input controls.

Ground-truth annotations for all 30 evaluation datasets were established through retrospective manual auditing of raw GEO metadata records and their associated peer-reviewed publications. The complete list of evaluated dataset accessions, expert-adjudicated classifications, and baseline retrieval results are detailed in Supplementary Table S1.

## Data availability

The GEOAgent integrated knowledge base, including the relational metadata database, vectorized knowledge cache, and neural re-ranking model weights, is available on Zenodo (https://zenodo.org/records/20138122). The standardized reference genomes and multi-omics indices configured for the bioStream pipeline are also deposited in a separate Zenodo record (https://zenodo.org/records/20045739).

## Code availability

A fully functional web implementation of the platform is directly accessible at https://geoagent.ccla.ac.cn. The open-source codebases for both the GEOAgent platform and the bioStream pipeline backend are available on GitHub: https://github.com/JiekaiLab/GEOAgent and https://github.com/JiekaiLab/bioStream. Additionally, the compiled VS Code extension artifact is distributed via the official release asset channel (https://github.com/JiekaiLab/GEOAgent/releases/tag/1.0.0).

## Author contributions

Yingying Zhao conceived and designed the study, performed data curation and database construction, developed the platforms and core bioinformatic pipelines, conducted the evaluation framework, and drafted and revised the manuscript. Quanyou Cai participated in project conceptualization, website deployment, benchmarking evaluation, and writing and reviewing the manuscript. Dongzhu Chen assisted in platform extension and interface development, and contributed to database construction, website deployment, and code testing. Jiekai Chen supervised the entire project and contributed to the critical review and editing of the manuscript. All authors have read and approved the final version of the manuscript.

## Competing interests

The authors declare no competing interests.

## Acknowledgements

This work was supported by the Strategic Priority Research Program of the Chinese Academy of Sciences (XDB1350000).The National Natural Science Foundation of China (32225012), Science and Technology Planning Project of Guangdong Province, China (2023B1212060050, 2023B1212120009), and Health@InnoHK Program launched by Innovation Technology Commission of the Hong Kong SAR, P. R. China.

## Notes

https://geoagent.ccla.ac.cn

https://github.com/JiekaiLab/GEOAgent

https://github.com/JiekaiLab/bioStream

### Competing Interest Statement

The authors have declared no competing interest.

