## Supplemental Note 1 for "GEOAgent: An AI-driven Autonomous Framework for Intelligent GEO Data Retrieval and Standardized Preprocessing"

**Supplementary Note 1.**

**Operational execution details for bioStream**

**1. Nextflow framework and reference standardization**

Once user-specified cohorts are curated and their corresponding Sequence Read Archive (SRA) download links are extracted, the framework dynamically compiles a standardized sample manifest (Samplesheet) and generates a single-line Nextflow execution command. This programmatic package is handed off to the user for localized, highly scalable data preprocessing via bioStream, an independent workflow architecture engineered natively in Nextflow (**Fig. 1, Step 7; Fig. 4**). By decoupling metadata curation from secondary resource-intensive computation, users can deploy the generated package directly onto their own high-performance computing (HPC) infrastructures. Analytical reproducibility and software portability are strictly enforced within the user environment by containerizing all software dependencies using Docker or Singularity images.

To eliminate reference-version mismatching across multi-modal integration, bioStream establishes a standardized reference genome architecture derived from a singular root provided by the 10x Genomics refdata-cellranger-arc ecosystem (**Fig. 4B**). The pipeline programmatically unpacks the baseline genomic assembly (genome.fa) and primary gene annotations (genes.gtf) to construct harmonized analytical indexes. This centralized setup generates the specific indexes required for STAR, RSEM, and Bowtie2, while synchronizing auxiliary configuration files—comprising ENCODE genomic blacklists (blacklist.bed) and ribosomal RNA intervals (rRNA.bed)—across concurrent execution paths.

**2. Multi-omics processing modules**

During execution, the bioStream pipeline automatically routes raw sequence data through specific processing paths based on the LLM-curated bioStream-Ready Samplesheet (**Fig. 4A**). This structured manifest encapsulates all upstream intelligence—including inferred omics modalities, organism origins, SRA retrieval links, and experimental pairing topologies—enabling fully automated, zero-intervention execution:

Data Acquisition and Quality Control: Raw SRA files are fetched via the SRA Toolkit, decompiled into FASTQ format using fasterq-dump, and profiled for raw sequence quality parameters via FastQC. Adaptive adapter and quality trimming are subsequently managed using Trim_Galore.

Bulk Transcriptomics: High-throughput RNA-seq reads are aligned to the reference genome using STAR. Transcripts are programmatically quantified via RSEM and featureCounts, while post-alignment quality metrics are systematically evaluated using RSeQC.

Epigenomics (ATAC-seq and ChIP-seq): Fragment alignment is executed via Bowtie2. The resulting BAM streams are filtered using Samtools and Picard to purge duplicate alignments and mitochondrial background noise. For ATAC-seq data, read coordinates are shifted by +4 bp on the forward strand and -5 bp on the reverse strand to adjust for transposase footprints (TN5_shift). Chromatin enrichments and protein-DNA binding sites are identified via MACS2 peak-calling, while continuous alignment densities are scaled and converted into bigWig tracks using deepTools (bamCoverage).

10x Genomics Single-Cell Assays: Raw FASTQ bundles are structured into technology-compliant folder topologies. Depending on the assay tag, the data are natively routed through the CellRanger suite utilizing cellranger count for scRNA-seq, cellranger-atac count for scATAC-seq, or cellranger-arc count for single-cell Multiome data to yield standardized expression matrices and accessibility fragments.

**3. Automated quality auditing and unified reporting**

To consolidate disparate tool-specific execution logs, bioStream integrates a dual-path summary reporting engine tailored to distinct assay modalities (**Fig. 4C**). For bulk sequencing tracks (RNA-seq, ATAC-seq, and ChIP-seq), standard alignment efficiencies, duplication rates, and trimming statistics—among other upstream metrics—are compiled via native MultiQC aggregation. Concurrently, for 10x Genomics workflows, the engine programmatically parses and embeds the interactive HTML web summaries generated directly by the CellRanger suite.

To incorporate specialized bulk epigenomic metrics, custom scripts calculate library saturation profiles via the Non-Redundant Fraction (NRF) and PCR Bottleneck Coefficient (PBC), monitor mitochondrial read ratios (Mito Ratio), and quantify signal specificity via the Fraction of Reads in Peaks (FRiP). These metrics are programmatically formatted as custom content modules following MultiQC specifications, allowing them to be seamlessly co-rendered with standard alignment metrics and localized metadata into a unified, evaluation-ready HTML dashboard and archival PDF report.
